# Selective inhibition of CA IX over CA XII in breast cancer cells using benzene sulfonamides: Disconnect between CA activity and growth inhibition

**DOI:** 10.1101/345298

**Authors:** Mam Y. Mboge, Zhijuan Chen, Alyssa Wolff, John V. Mathias, Chingkuang Tu, Kevin D. Brown, Murat Bozdag, Fabrizio Carta, Claudiu T. Supuran, Robert McKenna, Susan C. Frost

**Affiliations:** Department of Biochemistry and Molecular Biology, University of Florida, 1200 Newell Drive, Gainesville, FL 32610, USA; University of Florence, NEUROFARBA Department, Sezione di Farmaceutica e Nutraceutica, Via Ugo Schiff 6, 50019 Sesto Fiorentino (Florence), Italy

**Author notes:** **Corresponding author**, (SCF).

## Abstract

Carbonic anhydrases (CAs) have been linked to tumor progression, particularly membrane-bound CA isoform IX (CA IX). The role of CA IX in the context of breast cancer is to regulate the pH of the tumor microenvironment. In contrast to CA IX, expression of CA XII, specifically in breast cancer, is associated with better outcome despite performing the same catalytic function. In this study, we have structurally modeled the orientation of bound ureido-substituted benzene sulfonamides (USBs) within the active site of CA XII, in comparison to CA IX and cytosolic off-target CA II, to understand isoform specific inhibition. This has identified specific residues within the CA active site, which differ between isoforms that are important for inhibitor binding and isoform specificity. The ability of these sulfonamides to block CA IX activity in breast cancer cells is less effective than their ability to block activity of the recombinant protein (by one to two orders of magnitude depending on the inhibitor). The same is true for CA XII activity but now they are two to three orders of magnitude less effective. Thus, there is significantly greater specificity for CA IX activity over CA XII. While the inhibitors block cell growth, without inducing cell death, this again occurs at two orders of magnitude above the K_i_ values for inhibition of CA IX and CA XII activity in their respective cell types. Surprisingly, the USBs inhibited cell growth even in cells where CA IX and CA XII expression was ablated. Despite the potential for these sulfonamides as chemotherapeutic agents, these data suggest that we reconsider the role of CA activity on growth potentiation.

## Introduction

Breast cancer remains the second most diagnosed and one of the leading causes of cancer-related deaths among women in the United States. Gene expression patterns of dissected human breast tumors have provided distinct “molecular portraits” allowing classification into subtypes based solely on differences in these patterns [1]. These subtypes fall into three major groups: those that express estrogen (ER)/progesterone receptors (PR) (the luminal subtype), those that express HER2 (ERBB2+), and those that express neither (the basal-like, including the “triple negative” phenotype, TNBC). About 74% of breast cancer patients have ER-positive tumors [2]. For these patients, adjuvant therapy with tamoxifen (a selective estrogen receptor modulator) has contributed to the 30% decrease in breast cancer mortality in the past two decades, making it the most successful targeted cancer therapy to date [3]. Yet, about half of the tamoxifen-treated women will develop endocrine-resistant disease [4]. HER2 positive breast cancers are observed in about 15% of patients (some of which are also ER/PR positive), who in recent years has been successfully treated using a monoclonal antibody against the HER2 receptor. Triple negative breast cancer (TNBC) also accounts for about 15% of breast cancer patients, but is responsible for a disproportionate number of breast cancer deaths [5]. The higher mortality rate in women with TNBC is attributed to lack of targeted therapies coupled with the inability to treat metastatic disease [5-7]. That tumors develop resistance to radiation and/or chemotherapies complicates treatment.

Changes in the tumor microenvironment (TM), induced partly by hypoxia, underlie aggressive and resistant cancer phenotypes [8]. It is known that the hypoxic TM contributes to tumorigenic characteristics such as tumor cell proliferation, angiogenesis, cell motility, and invasiveness that translate to an overall poor prognosis in cancer patients [8, 9]. Hypoxia induces a global change in gene expression through stabilization of hypoxia inducible factor one alpha (HIF1α) [10]. Stabilization of this transcription factor leads to the upregulation of genes important for cell survival [10]. Hypoxia-induced upregulation of genes in glycolysis cause elevated glucose uptake, which is a hallmark for actively dividing cells and correlates closely with transformation [11-13]. Increased glycolysis results in high rates of extracellular acidification (pH_e_) through the export of H^+^ in an attempt to maintain a more basic/neutral intracellular pH (pH_i_) [14-16]. This increase in H^+^ production and export to the extracellular space induces apoptosis in normal cells surrounding tumors [17]. In contrast, by mechanisms not understood, cancer cells establish as new “set-point” which allows them to tolerate low pH_e_ [8, 17-19]. The stress of the TM also induces resistance to therapeutic drugs, preventing uptake as pH gradients change across the membrane [12]. This, in turn, induces the counter transport of drugs reducing concentrations inside cells and thus effectiveness. This is true not only for TNBC patients treated with cytotoxic drugs but also estrogen receptor (ER)-and HER2-positive patients treated with targeted therapies [3].

Important modulators of the hypoxic TM include membrane-bound carbonic anhydrases (CAs), which are zinc metalloenzymes that catalyze the reversible hydration of CO_2_ [16, 20-26]. In breast cancer, carbonic anhydrase IX (CA IX) expression is upregulated in hypoxic tissue and is considered a prognostic marker for the TNBC phenotype [27-30]. In normal cells, CA IX expression is limited primarily to gut epithelium, but is upregulated in many hypoxic tumors including breast cancer [28, 31-43]. CA IX is not only a marker for hypoxia but may also serve as a therapeutic target for the treatment of aggressive breast cancer. CA XII expression, on the other hand, is not regulated by hypoxia and is an indicator of better patient outcome in several types of cancers, including breast cancer [44]. Its expression is under the control of the ER and associated with ER-positive breast carcinomas [44-47]. CA XII is also expressed more abundantly across tissue types, including normal breast tissue [41, 48].

The role of these carbonic anhydrases in cancer progression are under intense investigation, particularly that of CA IX. CA IX was originally identified as a subunit of the MaTu protein and a short time later characterized as a novel member of the carbonic anhydrase family [49, 50]. While it is normally localize to gut epithelium, it is strongly upregulated in tumors, generally in more aggressive phenotypes [51, 52]. CA IX is a transmembrane protein, organized as a dimer, with the catalytic domain facing the extracellular space. This enzyme catalyzes the reversible hydration of CO_2_ forming a bicarbonate ion and a proton, in the presence of water. The pericellular space, with which CA IX associates, is rich in metabolic acids such as lactate, protons, and CO_2_. It has been postulated that CA IX cooperates with bicarbonate transporters by providing bicarbonate for intracellular alkalinization at the same time that it acidifies the tumor microenvironment [53]. Data supporting this hypothesis have been provided through studies in cell lines and cancer cell-derived spheroids [34, 54, 55]. However, CA IX, like other members of the family, is a reversible enzyme. We, and others, have hypothesized that CA IX may stabilize extracellular pH at a value that favors cancer cells growth and proliferation [27, 56]. New data from studies with CA IX expressing xenografts support this notion that CA IX acts as an extracellular pH-stat [57]. This adds to the therapeutic rationale for pharmacological interference with CA IX activity.

Sulfonamides are the most studied class of CA inhibitors (CAIs) with several compounds in clinical use for the treatment of diseases including glaucoma, altitude sickness, and epileptic seizures [25, 58-62]. Although these sulfonamides are potent CAIs, most lack isoform selectivity and do not specifically target the tumor-associated isoforms CA IX and CA XII [60]. This lack of specificity is attributable to the highly conserved residues within the active site of CAs [63]. These similarities have challenged the development of isoform-specific inhibitors, causing competition between isoforms, which reduce efficacy. This is especially true in the case of CA II. This isoform has the widest tissue distribution and is abundantly expressed in red blood cells (representing 5% of all proteins) [64, 65]. As most drugs are delivered systemically and are membrane permeable, it is likely that CA II will sequester CA inhibitors reducing circulating concentrations limiting exposure within tumors. That said, none of the clinical CAIs have been used for the treatment of cancer. Ureido-substituted benzene sulfonamides (USBs) are potent and selective inhibitors of recombinant CA IX and CA XII activity [66-68]. X-ray crystallographic studies recently provided insight into the mechanism of USB-mediated selective inhibition of CA IX over CA II [66]. Those studies showed that the USBs interact more favorably with residues in the active site of CA IX versus CA II [66]. These inhibitors also showed both anti-proliferative and anti-metastatic properties in preclinical models [29, 68]. One of the USBs (SLC-0111) recently completed phase I clinical trials and is scheduled to begin phase II trials for the treatment of solid tumors that over express CA IX (clinical trials.gov, NCT02215850).

The goal of this study was to distinguish the mechanistic effects of USB-mediated inhibition of CA IX and CA XII in the context of breast cancer cells. To accomplish this, three USBs (including SLC-0111) were studied, using *in-silico* modeling to understand binding mechanisms and their efficacy was tested in cell lines representing TNBC and ER-positive breast cancers. With this approach, differences in residues within the active sites of CA IX and CA XII indicated that these drive isoform-specific interactions affecting affinity of the USBs. The USB compounds were also shown to inhibit CA activity with surprising specificity for CA IX in TNBC cells, relative to CA XII in ER-positive cells given similar K_i_ values for purified protein. This same specificity appears to drive USB-mediated growth inhibition. Yet the K_i_ values for inhibiting catalytic activity in CA IX and CA XII in cells is two and three orders, respectively, less efficacious than would be expected based on activity studies using purified protein and binding kinetics based on structures. In addition, growth inhibition requires even higher concentrations of inhibitors than that needed for inhibition of activity in cells. Further, cells that lack expression of CA IX or CA XII are still sensitive to USB-mediated inhibition of cell growth. Despite the excellent inhibition of the USBs against purified CA IX and CA XII, and the specificity for CA IX over CA XII in cells, we must consider the possibility that CA activity may not be the target for sulfonamide action with respect to cell growth inhibition.

## Materials and methods

Ureido-substituted benzene sulfonamide (USB) inhibitors were prepared in dimethyl sulfoxide (DMSO) at stock concentrations of 100 mM and further diluted to appropriate concentrations in cell culture medium before use. Control experiments contain DMSO only. CA II antibody was obtained from Novus Biological (Littleton, CO), CA IX (M75) antibody was a gift from Egbert Oosterwijk, CA XII and GAPDH antibodies were purchased from R&D systems (Minneapolis, MN) and Cell Signaling Technology (Beverly, MA), respectively.

### *In Silico* Modeling

X-ray crystallography structures of USB compounds in complex with purified CA II and a CA IX-mimic (analogous site directed mutagenesis of residues in the active site of CA II to resemble CA IX) were previously obtained and published elsewhere [66], but shown for comparison to the modeled CA XII complex in the present study (Figure 1). Each of the USB compounds were superimposed into CA XII structures (PDB ID: 1JCZ) based on the crystal structures of the compounds using Coot [69, 70] and adjusted based on steric hindrance constraints. These structures were also energy minimized using the Chimera Minimize structure feature (data not shown) to ensure that these modeled compounds were at their energy minima.

### Cell Culture

MCF 10A, UFH-001, and T47D cells were maintained as previously described [58]. For hypoxia treatment, cells were exposed to 1% O_2_, 5% CO_2_ and balanced N_2_ for 16 h using a Billups Rothenberg Metabolic Chamber. UFH-001 cells are a newly characterized line that exhibits the triple negative (TNBC) phenotype [71, 72]. These cells do not express ER, the progesterone receptor (PR), or HER2. However, they do express EGFR, also typical of the TNBC phenotype. In addition, these cells express CA IX but not CA XII. The T47D cells are a luminal ER-positive line that expresses CA XII but not CA IX. The MCF 10A line is used as a control and expresses CA IX only under hypoxic conditions. All cell lines were authenticated prior to conducting experiments.

### CA IX deletion by CRISPR/Cas9

Ablation of CA IX in UFH-001 cells has been described previously [72]. pLentiCRISPR v2 was a gift from Dr. Feng Zhang (Addgene, Cambridge, MA) and was used to knockout CA IX in UFH-001 cells. Briefly, guide RNA sequences within the first coding exon were identified downstream of the CA IX translational start site using an online design tool (crispr.cos.uni-heidelberg.de) and three non-overlapping gRNAs were chosen within this region of the *CA9* gene. Complementary double-stranded oligonucleotides were cloned into BsmBI digested pLentiCRISPR v2, recombinant clones identified by restriction analysis and confirmed by automated Sanger sequencing. Lentivirus was formed from these CA IX sgRNA plasmids and empty lentiCRISPR v2 plasmid as previously outlined [72]. UFH-001 cells were transduced and selected with puromycin (2 µg/mL) for 3 weeks. Stably transduced cells were harvested, and CA IX depletion was confirmed by western blotting.

### Knockdown of CA XII by shRNA lentiviral particles

Knockdown of CA XII expression in T47D cells has also been previously described [72]. Briefly, cells were transfected with short hairpin RNA (shRNA) lentiviral particles obtained from Thermo Fisher Scientific (Waltham, MA) against CA XII. Cells were seeded at 5×10^4^ cells/well in 24-well plates and grown for 24 h. Then, cells were infected with lentivirus in serum-free medium for 6 h. The cells were further incubated with normal growth medium for 24 h. GFP expression was monitored to confirm the efficiency of transduction. Stable cells were established by puromycin (2 µg/mL) selection (Sigma Aldrich, St. Louis, MO). CA XII knockdown was confirmed by western blotting.

### Western Blot Analysis

Western blot analysis was performed, as previously described [28]. Briefly, cells were collected after drug treatments for 48 h and washed 3x with ice-cold phosphate buffered saline (PBS) and lysed in RIPA buffer (containing 1% orthovanadate, 25 mM NaF) supplemented with protease inhibitor for 15 min on ice. Lysates were collected and clarified by centrifugation at 55,000 rpm for 60 min at 4°C. Clarified supernatants were collected and stored at-20°C. Protein concentration was determined using the Markwell modification of the Lowry procedure. Equal amounts of cell lysate (30 μg) were subjected to SDS-polyacrylamide electrophoresis, prepared as described by Laemmli et al. [73]. Samples were electrotransferred from SDS-PAGE gels to nitrocellulose membranes in transfer buffer at 200 mA for 2 h at 4°C. After incubation with selective primary and secondary antibodies, proteins were visualized using ECL reagent according to the manufacturer’s directions (GE Healthcare, Wauwatosa, WI). Images were scanned and cropped using Adobe Photoshop version 11.

### Membrane Inlet for Mass Spectrometry

CA catalysis was measured by the exchange of ^18^O from species of ^13^CO_2_ into water determined by membrane inlet mass spectrometry (MIMS) [74, 75]. The application of this method in demonstrating exofacial CA activity in breast cancer cells has been described previously [27] and more recently in Chen et al. [72]. To decrease inaccuracies arising from ^12^C-containing CO_2_ in cell preparations, ^13^C-and ^18^O-enriched CO_2_/HCO_3_ were used and the rate of depletion of ^18^O in ^13^C-containing CO_2_ measured. Specifically, the atom fraction of ^18^O in extracellular CO_2_ was determined using peak heights from the mass spectrometer representing CO_2_ [1/2 (47) + (49)] /[(45) + (47) + (49)] where the numbers in parentheses represent the peak heights of the corresponding masses of specific species of isotopic CO_2_. The MIMS assay actually measures CA activity in the dehydration direction, because only isotopic variations in CO_2_ are measured by the mass spec (although activity in both directions is required for the decrease in the isotopic distribution of ^18^O in CO_2_). A membrane barrier (to the mass spec) prevents bicarbonate concentration from being measured by the mass spec. When added to the reaction vessels, the cells are exposed to an equilibrated solution of^13^C^18^O_2_:HC^18^O_3_^-^ at a ratio of about 5%: 95%, respectively (25 mM, total). The total amount of CO_2_ does not change over the course of the experiment, just the isotopic enrichment. Mass spectra were obtained on an Extrel EXM-200 mass spectrometer using electron ionization (70 eV) at an emission current of a 1 mA. First order rate constants (k) were determined for CA activity. These were calculated from the change in atom fraction enrichment (^18^O in CO_2_) using: A = A_0_ e^-kt^, where (A) is the fraction of enrichment at a specific end point, and (A_0_) is the starting enrichment at a specific point.

Cells were collected from culture plates using cell release buffer (Gibco, Waltham, MA), washed with bicarbonate-free DMEM buffered containing 25 mM Hepes (pH 7.4), and counted. Cells (5.0 × 10^5^ cells/mL, unless otherwise stated) were added to a reaction vessel containing 2 mL of Hepes-buffered, bicarbonate-free DMEM at 16° C in which ^18^O-enriched ^13^CO^2^ /H^13^CO_3_ at 25 mM total ^13^CO_2_ species was dissolved. Catalysis by CA, in the presence or absence of CA inhibitors, was measured after addition of cells (exposed to normoxic or hypoxic conditions) by the exchange of ^18^O species from species of ^13^CO_2_ into water determined by measuring CO_2_ mass by MIMS as described above. This exchange is a specific measure of CA activity during the time frame of the assay. A general example of this assay is provided in Supporting Information (S1 Fig) showing the effect of two classical sulfonamide based inhibitors on CA activity in UFH-001 and T47D cells, which is further discussed in the results.

Ki values were calculated using the following equation:

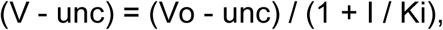

where (V) is the rate at a specific inhibitor concentration, (unc) is the uncatalyzed rate, (Vo) is rate in the absence of inhibitor, and (I) is inhibitor concentration. For tight binding inhibitors, IC_50_ = [enzyme]/2 + Ki (app). However, when the K_i_ >> greater than the enzyme concentration, IC_50_ ≃ K_i_. In our experiments, enzyme concentration in cells is in the low nM range, based on titrations with purified CA II. The K_i_ values that we calculate are much higher than [enzyme] for either UFH-001 or T47 D cells, despite the difference in enzyme concentration between the two cell lines.

### Cell Growth Assay

Cell growth in the presence or absence of compounds was analyzed using 3-[4,5-dimethylthiazol-2-yl]-2,5-diphenyltetrazolium bromide (MTT assays). Briefly, cells were seeded in 96-well plates and treated with different concentrations of compounds after 24 h. The cells were then incubated under normoxic or hypoxic conditions under the different treatment conditions for 48 h and/or 96 h. The medium was replaced every two days. MTT reagent (10% v/v of 5 mg/mL) was added to the cells after each time point and the cells were incubated for an additional 4 h. The medium was removed without disturbing the cells and DMSO was added to each well. The DMSO dissolved cells were incubated while shaking for 15 min in the dark. Absorbance was measured at 570 nm using an Epoch microplate reader (Biotek, Winooski, VT). IC_50_ values (inhibitor concentration where the response is reduced by half) were calculated using Prism 7.0c for Macs. Non-linear regression analysis was performed on % dose response with log plots interpolating the value at 50% from the least squares fit of the growth curves.

### Lactate Dehydrogenase Release Assay

Cytotoxicity was estimated using the release of lactate dehydrogenase (LDH) activity (Sigma-Aldrich, St. Louis, MO). Cells were plated in 96-well plates. USB compounds were added the next day. After 48 h, medium was collected from each well to measure the amount of released LDH activity. The amount of LDH released from each sample was measured at room temperature at 450 nm using a BioTek Epoch microplate reader. LDH activity is reported as nmol/min/mL of medium.

### Caspase-3 Activity Assay

Caspase-3 activity assay was performed using an Apo-alert colorimetric caspase assay kit (BioVision, Milpitas, CA). Cells were plated onto 10 cm dishes and treated with compounds 24 h later. 48 h after various treatments, cells were collected, and lysates prepared. Protein concentration was determined, and equal volumes of lysates were used for caspase-3 activity assay, measured at 405 nm in a microtiter plate. The reader detected chromophore p-nitroaniline (pNA) after its cleavage by caspase-3 from the labeled caspase-3 specific substrate, DEVD-pNA. These data are presented as pmol of pNA per µg of cell lysate per hour of incubation. Staurosporine (Sigma-Aldrich, St. Louis, MO) was used as a positive control.

### Analysis of Cell Cycle Progression

Cells were seeded in 24-well plates. After 24 h, USB compounds were added to the respective wells and incubated for 48 h. Cells were released with trypsin, harvested, fixed with 500 µL of 50% ethanol in tubes and collected by centrifugation at 13,000 rpm for 5 min. Cell pellets were re-suspended in 500 μL. Propidium Iodine (10 µg/mL) containing 300 µg/mL RNase (Invitrogen, Waltham, MA). Then, cells were incubated at room temperature in the dark for 15 min. Cell cycle distribution data in Figure 8C-8E (gated for live, diploid cells only) was analyzed using the FCS Express Version 5 from De Novo Software (Glendale, CA) from data obtained using FACS caliber (Becton Dickson, CA). The representative images in the text (Fig 8A and 8B) were generated by ModFit LT.

### Cell Migration and Invasion Assays

Serum starved cells were plated at a density of 50,000 cells in 300 µL per insert in 24-well cell migration and invasion plates (Cell BioLabs, San Diego, CA). The cells were allowed to migrate (24 h) or invade (48 h) from the insert containing serum free medium, with or without USB compounds, towards the well that had medium containing 10% FBS. Fixing and staining cells terminated the assay. Images were then collected and analyzed. All cell lines were maintained at 37°C in humidified air with 5% CO_2_ for the duration of the experiment.

### Spheroid Formation Assay

UFH-001, UFH-001 CA IX KO, T47D, and T47D CA XII KO cells (10,000 cells in a total volume of 100 µL) were plated into 96-well, low attachment microplates (Corning, NY). Cells were incubated up to 96 h in 5% CO_2_ at 37°C. Spheroids were imaged at specific intervals using the EVOS microscope system (Thermo Fisher Scientific, Waltham, MA).

### Statistical Analysis

Data are presented as means ± SEM. Differences between the treatment groups were analyzed using Student’s t-test in GraphPad Prism, version 7.0a: *p*-values < 0.05 was considered statistically significant.

## Results

### Isoform selective binding and interaction of USB compounds to CA IX and CA XII

Cytosolic CA II exists as a monomer (Fig 1A), while both membrane-bound CA IX and CA XII have been shown to form dimers. The catalytic domains of these dimers are shown in Fig 1C and 1E. The structural formulae of the USB inhibitors used in this study and their respective K_i_ values for recombinant CA II, IX and XII were determined elsewhere and provided in Table 1 [66-68]. These previously published data show that two of the three USB compounds (U-CH_3_ and U-F, also called SLC-0111) selectively bind to and inhibit the activity of the tumor-associated isoforms CA IX and XII relative to the off-target CA II [66-68]. Isoform selectivity towards recombinant CA IX and CA XII was evident in the differences in K_i_ values for these isoforms relative to CA II, obtained using Applied Photophysics stopped-flow kinetics of purified recombinant proteins. Isoform selectivity is important *in vivo*, where inhibition of CA II (one of the predominant isoforms in red blood cells) might cause unintended consequences and sequester inhibitor reducing the concentration within the tumor.

**Table 1.**
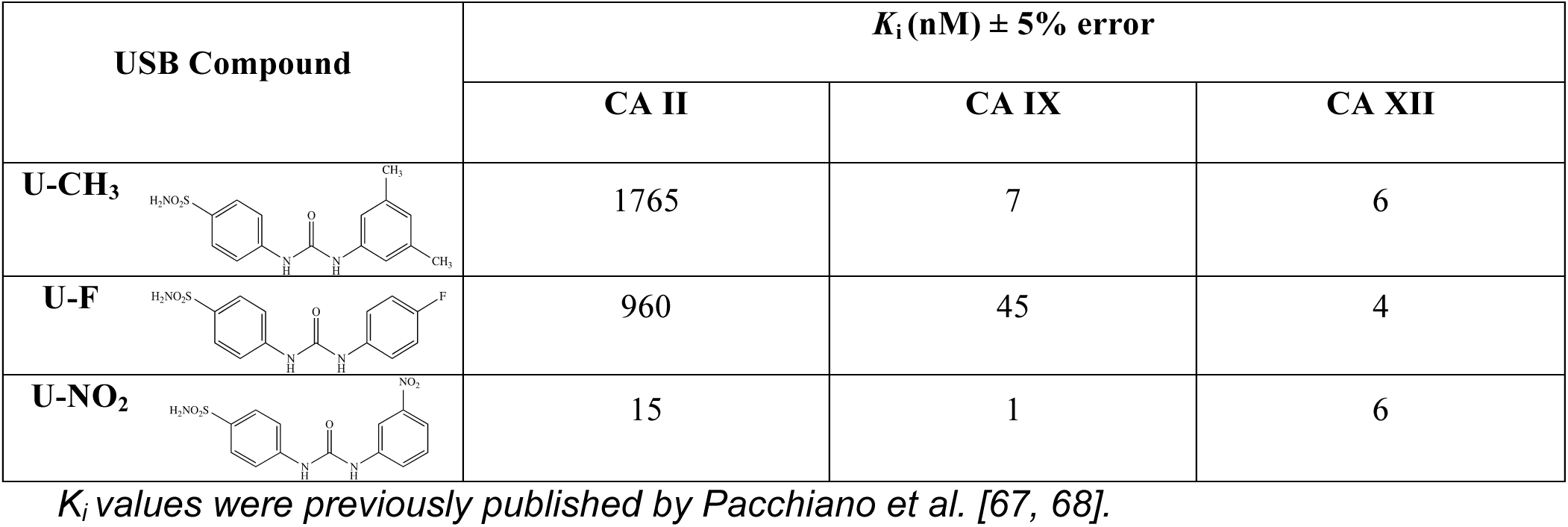
Inhibition constant (K_i_) values for recombinant human CA II, IX, and XII with ureido-substituted benzene sulfonamide (USB) compounds **U-CH_3_, U-F,** and **U-NO_2_** obtained from Applied Photophysics stopped-flow kinetics experiments.

| USB Compound | $K_i$ (nM) ± 5% error | | |
| --- | --- | --- | --- |
|  | CA II | CA IX | CA XII |
| **U-CH<sub>3</sub>** | 1765 | 7 | 6 |
| **U-F** | 960 | 45 | 4 |
| **U-NO<sub>2</sub>** | 15 | 1 | 6 |

$K_i$ values were previously published by Pacchiano et al. [67, 68].

To investigate the underlying mechanism of binding, we previously determined the crystallographic structures of all three compounds bound to CA II (Fig 1B) and the CA IX-mimic (Fig 1D) [66]. As expected, each compound was shown to directly interact with the zinc ion, via the sulfonamide moiety [66]. The extended tail moieties of the USB compounds also interacted with residues, between 10-15 Å away from the zinc, favoring the hydrophobic region of the active site (Fig 1B and 1D). This region is located close to the entrance of the active site and has been termed the “selective pocket” [76, 77]. This is because it has a clustering of several amino acid residues that are different among CA isoforms (Table 2). For example, residue 131, located within the selective pocket, is different among the isoforms. In CA II, this residue is a phenylalanine (CA II residue numbering will be used throughout this manuscript). Its bulky side chain causes steric hindrance and therefore induces conformation changes in the tail groups of the bound compounds (Fig 1B, Table 1). In CA IX, residue 131 is a valine, which allows for more favorable binding and therefore greater isoform selectivity towards CA IX compared to CA II (Fig 1D, Table 1). We believe that the differences in active site residues, located within the selective pocket, are responsible for the observed differences in K_i_ values for bound compounds (Table 1). By comparison, the classical CA inhibitor, acetazolamide, can only interact with the active site zinc ion and with residues within 10 Å of the zinc, because of its shorter tail moiety [78]. Residues in this region of the active site are among the most conserved between isoforms [63].

To compare the CA II and CA IX structural data with that of CA XII, we have now performed *in silico* modeling of the three USB compounds (U-CH_3_, U-F and U-NO_2_) into the active site of CA XII (Fig 1F). Residue 131 in CA XII is an alanine (Table 2), which like valine (in CA IX) has a small hydrophobic side chain that results in favorable binding and selectivity towards this isoform (Table 1). In addition, CA XII has a serine (a polar uncharged side chain) at position 132, which is also located close to the entrance of the active site. As such, this residue also causes steric hindrance that induces conformational changes in all three sulfonamides, enhancing binding and inhibition (Table 1). Additionally, serine 132 and also serine 135 (both charged residues and also unique to CA XII) create a narrow hydrophobic region within the CA XII active site (Fig 1F). This causes an energetically favorable change, especially, in the conformation of compound U-F (relative to CA II and CA IX) within the active site of CA XII. These residues also prevent steric clashes with residue 131 and facilitate compound interactions in the narrow hydrophobic region within the selective pocket. These interactions and the resulting conformational change in U-F will be specific to CA XII, because neither CA II nor CA IX has serine residues at positions 132 or 135 (Fig 1 and Table 2).

**Fig 1.**
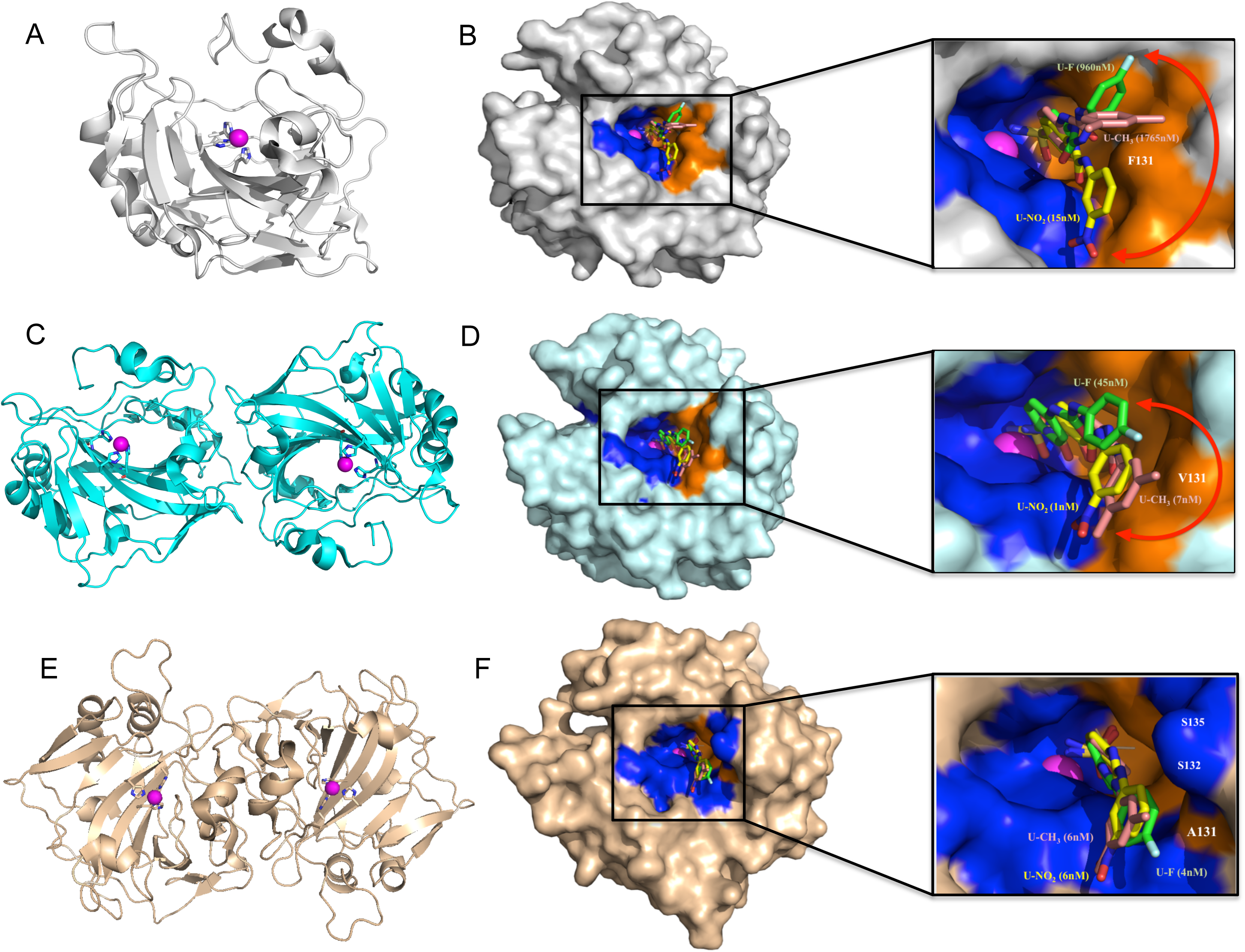
USBs bound in the active site of CAs. Panel A. Ribbon diagram of monomeric CA isoform II (gray). Panel C. Ribbon diagram of the catalytic domains of dimeric CA isoform IX (cyan). Panel E. Ribbon diagram of the catalytic domains of dimeric CA isoform XII (wheat). Surface representation of compounds U-CH_3_ (pink), U-F (green) and U-NO_2_ (yellow) in complex with monomers of CA II (Panel B), the CA IX-mimic (Panel D), and modeled into the active site of CA XII (Panel F). Catalytic zinc (magenta sphere), hydrophilic (blue) and hydrophobic (orange) residues are as shown. Red double-headed arrows indicate isoform specificity relative to residue 131 (labeled in white). These arrows also show flexibility in tail conformations seen in CA II and CA IX but not in CA XII. Previously published K_i_ values of each compound bound to purified CA II, CA IX and CA XII are noted next to the inhibitor name [67, 68]. Figures were designed using PyMol.

**Table 2.** Resides within the active site of human CA II, CA IX and CA XII that differ among all three isoforms and make up the selective pocket (*CA II numbering*)

| Residues in Selective Region | CA II | CA IX | CA XII |
| --- | --- | --- | --- |
| **67** | Asn | Gln | Lys |
| **91** | Ile | Leu | Thr |
| **131** | Phe | Val | Ala |
| **132** | Gly | Ala | Ser |
| **135** | Val | Leu | Ser |

Information adapted from Pinard et al. [63]

### Effect of USB inhibitors on CA IX and CA XII activity in breast cancer cells

One of the goals of this study was to determine if these high affinity inhibitors (for CA IX and CA XII) were equally effective in the context of the cellular environment. To determine the effect of USB compounds on CA activity in breast cancer cells, cell lines that only express CA IX or CA XII at the cell surface were selected. Based on previous work, two lines were selected based on their strong expression of CA IX or CA XII. The T47D line is a well-studied ER-positive, luminal line. UFH-001 cells represent a new line, with a triple negative phenotype, arising from MCF 10A cells. MCF 10A cells were derived from a patient with fibrocystic disease, which spontaneously immortalized in cell culture [79, 80]. These are considered by many as the control line for breast cancer cells, although these cells map with other lines that have the basal/triple negative phenotype [81]. The UFH-001 cells are an especially aggressive that, unlike MCF 10A cells, form tumors in nude mice. Characterization of the UFH-001 cells is described in two recent publications [71, 72]. When compared to control MCF 10A cells, UFH-001 cells show greater expression of CA IX protein (Fig 2A) and mRNA (S2A Fig) under normoxic conditions. However, under hypoxic conditions both MCF 10A and UFH-001 cells upregulate CA IX expression (Fig 2A). It can also be noted that there is a difference in the migration patterns of CA IX between the two cell lines. In UFH-001 cells, CA IX migrates as a doublet with molecular weights of 54 and 58 kDa, both of which are glycosylated [82]. However, in MCF 10A cells, CA IX migrates as a singlet with a molecular weight slightly lower than 58 kDa. The observed differences in molecular weight may result from differences or deletions in primary sequence. CA XII expression is limited to the T47D cells (Fig 2B and S2B Fig) and is independent of hypoxia (Fig 2B). Expression of cytosolic CA II protein is only observed in UFH-001 cells and is also not sensitive to hypoxia (Fig 2A). The upregulation of CA IX in hypoxic UFH-001 cells underlies the increase in cell surface CA activity in hypoxic versus normoxic cells (Fig 2C) [27]. That there is no change in CA activity in T47D cells in response to hypoxia relative to normoxia also correlates with the lack of change in CA XII expression (Fig 2D).

**Fig 2.**
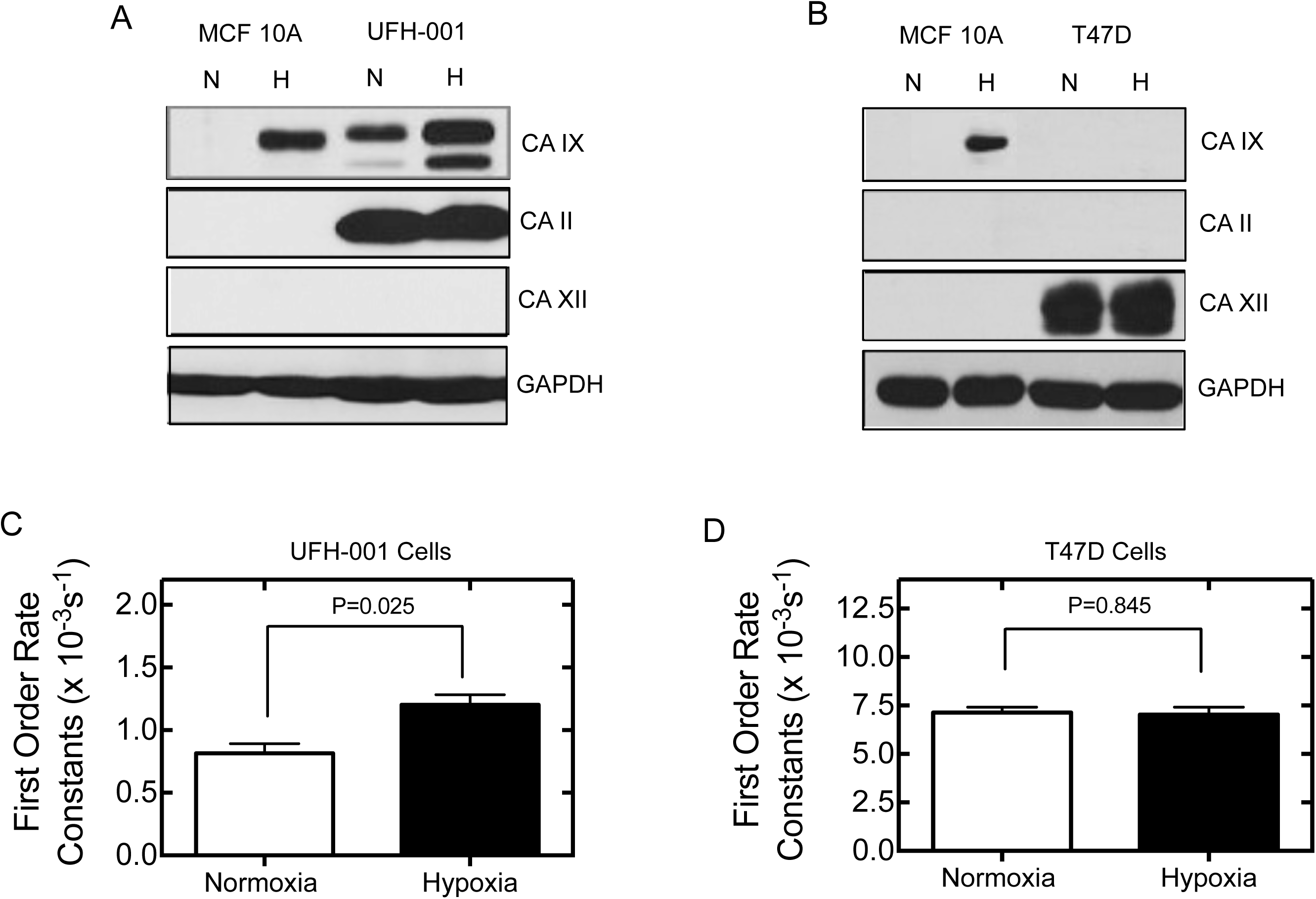
CA expression and activity in breast cell lines. Panel A. Protein expression in a normal immortalized basal type breast cell line (MCF 10A) and a triple negative breast cancer (TNBC) cell line (UFH-001), using western blot analysis (under normoxic (N) or hypoxic conditions (H) for 16 h, respectively). Panel B. Immunoblots for CAs II, IX and XII protein expression in MCF 10A and the luminal ER positive breast cancer cell line (T47D). Panels C and D. UFH-001 and T47D cells were grown for 3 days at which point they were exposed to normoxia or 16 h of hypoxia and the cells assayed for CA IX and XII activity, respectively, using the MIMS assay. First order rate constants for CA activity in UFH-001 cells (Panel A) and T47D cells (Panel B), as described in the methods, are shown. These data represent three independent experiments and are reported as the mean ± SEM.

The inhibitory capacity of USB compounds U-CH_3_, U-F, and U-NO_2_ on CA IX and CA XII activity in UFH-001 and T47D cells, respectfully, were studied under normoxic and hypoxic conditions, but only the representative normoxic plots are shown (Fig 3). Because UFH-001 cells express both CA II and CA IX, the progress curves that follow CA activity are biphasic on a semi log plot (Fig 3A-3C). The first phase (phase I, 20-40 sec after addition of cells) reflects CA II activity. The second phase (phase II, 100-400 sec) reflects CA IX activity. Phase I represents the rapid diffusion of ^13^C^18^O_2_ into cells where CA II catalytic cycles deplete ^18^O from H^13^C^18^O_3_ followed by efflux of ^13^CO_2_ species from the cell. The concentration of CO_2_ and the isotopic forms of CO_2_ are measured by the mass spec. There is little change in the slope of this phase between normoxic and hypoxic cells, although the length of time in this phase is shortened (Fig 3A-3C). Phase II represents the hydration-dehydration activity of CO_2_/HCO_3_^-^ outside of the cell representing CA IX activity. The first order rate constant of phase II is greater for hypoxic cells compared with normoxic cells (Fig 2C). This indicates a higher rate of CO_2_ hydration/dehydration on the extracellular face of hypoxic UFH-001 cells, which we associate with enhanced expression of exofacial CA IX (Fig 2A and 2C).

The results in Fig 3 also show that USB compounds have limited effects on phase I in the UFH-001 progress curves under either normoxic (Fig 3A-3C) or hypoxic conditions (data not shown). This means that during the time course of these experiments, the inhibitors did not permeate the membrane to inhibit CA II except at high concentrations of U-F (SLC-0111) which was observed as an increase in the atomic fraction of ^18^O in ^13^CO_2_ (starting at ∼150 seconds, Fig 3B). All three CA inhibitors had significant effects on phase II of the progress curve indicative of a decrease in CA IX activity (Fig 3A-3C). In other words, the slopes of the phase II progress curve (used to calculate first order rate constants in Fig 2) decline in the presence of the inhibitors. Based on K_i_ values in normoxic and hypoxic UFH-001 cells, the most potent inhibitor was U-F (SLC-0111) and the least inhibitory was U-NO_2_ (Fig 3G). Hypoxia did not significantly affect the efficacy of any of the USB compounds. These plots are representative of multiple experiments but show a limited number of inhibitor concentrations. The type of experiments that are used to actually calculate K_i_ values are shown in S3 Fig where an extensive range of concentrations (for U-NO_2_ in these cases) was utilized. In addition, we have included the effect of two classical sulfonamide inhibitors on CA activity in UFH-001 and T47D cells, that differ in their cell permeability (S1 Fig). Acetazolamide (ACZ) is impermeant during the time course of the experiment and is expected to block only exofacial CA activity, like CA IX and CA XII. Ethoxzolamide (EZA) is permeant, and quickly inhibits both exofacial and intracellular CAs. Activity in UFH-001 cells (S1A Fig), is, once again, observed as a biphasic progress curve. The presence of ACZ blocks only exofacial CA activity which confirms that the second phase of the progress curve (from about 200 to 600 sec) represents CA IX activity. The activity in the presence of EZA represents the spontaneous (non-catalytic) interconversion of CO_2_ and HCO_3_^-^, which is essentially an extension of the activity before addition of cells. First order rate constants are shown for each of the progress curves (S1A Fig). Finally, we present data on CA activity in cells in which CA IX is ablated (Fig 4A). Here, we show that CA activity is reduced in CA IX KO cells relative to the EV (hypoxic) controls. However, inhibition by the impermeant sulfonamide, N-3500, is better at blocking CA activity than is CA IX ablation. It is possible that there are other exofacial CA’s present on the surface of UFH-001 cells. We have only tested for the expression of CA XIV, which is not detected under conditions in which we observe CA IX. While microarray data (S2A Fig) show CA XIV mRNA expression in UFH-001 cells, earlier northern blot analysis revealed no CA XIV mRNA [23]. Furthermore, no difference in CA XIV mRNA expression was observed between MCF 10A and UFH-001 cells, under normoxic conditions (S2A Fig). As far as we are aware, no one has shown positive expression of either CA XIV or CA IV (the other membrane bound CA) at the protein level in breast cancer cells.

T47D cells, which express membrane-bound CA XII (Fig 2B), exhibit single-phase progress curves (Fig 3D-3F), because they lack CA II. This activity specifically reflects exofacial CA XII activity. This activity was not sensitive to hypoxia (Fig 2D), which is consistent with lack of CA XII induction by hypoxia (Fig 2B). Like CA IX, USB compounds inhibit CA XII activity (Fig 3D-3F). Again, the slope of the progress curve decreases in the presence of inhibitor. However, there is a striking difference in efficacy that is reflected in a significant increase in the K_i_ values obtained in T47D cells versus UFH-001 cells (Fig 3G). Again, an extensive range of concentrations are used to calculate K_i_ and shown for one of the inhibitors (U-NO_2_) in S3 Fig. Unlike UFH-001 cells, there were only slight differences in inhibitory effects of the different USB’s in T47D cells, which again was not affected by normoxic or hypoxic conditions (Fig 3G). S1B Figure shows the progress curves in the presence of ACZ and EZA. Catalysis in the absence of inhibitors is linear. Inhibition by ACZ generates a progress curve that resembles that of USB compounds. The activity in the presence of EZA once again represents the spontaneous (non-catalytic) interconversion of CO_2_ and HCO_3_^-^. These experiments were performed at 16°C, as were all of the MIMS experiments in this manuscript, so can be directly compared. CA XII knockdown significantly reduced CA activity in T47D cells relative to EV controls (Fig 4B). Inhibition of CA activity with N-3500 is only slightly more effective than the knockdown of CA XII expression.

**Fig 3.**
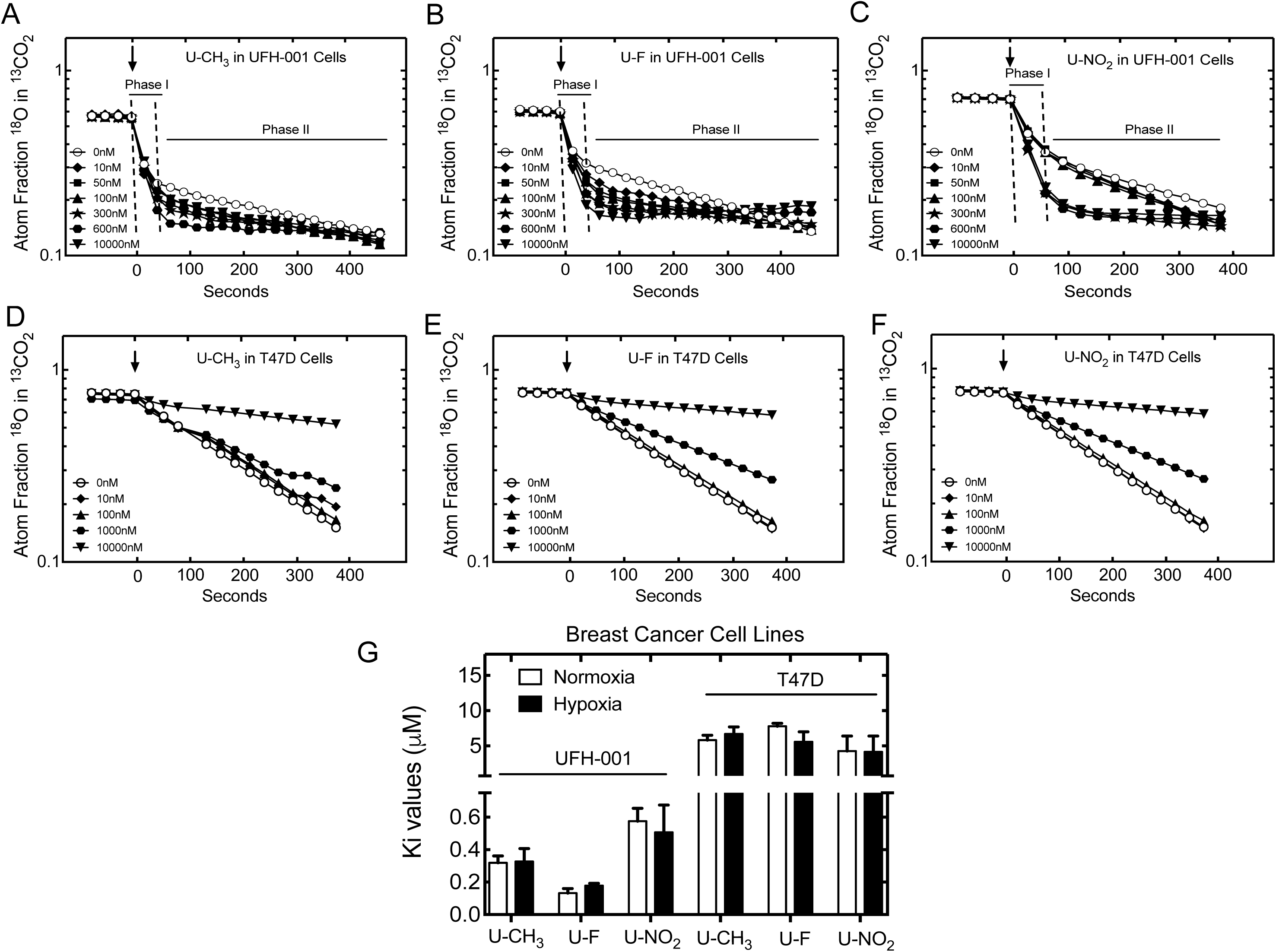
Effects of USBs on CA IX and CA XII activity in breast cancer cell lines. Panels A-C. UFH-001 cells were grown for 4 days under normoxic conditions. CA IX activity (0.5 × 10^6^ cells/mL, unless otherwise indicated) was assayed using MIMS in the presence or absence of U-CH_3_ (Panel A), U-F (Panel B) or U-NO_2_ (Panel C). Panels D-F. T47D cells prepared similarly to UFH-001 cells, but cultured for 6 days, were also assayed for CA XII activity (0.5 × 10^6^ cells/mL) in the absence or presence of the same USB based inhibitors: U-CH_3_ (Panel D), U-F (Panel E) or U-NO_2_ (Panel F). Atom fractions of ^18^O in CO_2_ were collected continuously. For ease of illustration, data points at 25-s intervals are shown. Phase I in UFH-001 cells indicate CA II activity and Phase II indicate CA IX activity. In T47D cells, the progress curves are linear and representative of CA XII activity. Arrows indicate time point at which cells were added. Panel G. K_i_ values of each compound in UFH-001 and T47D cells, under normoxic or hypoxic (16 h) conditions, are shown. Data are representative of the average of triplicate experiments ± SEM.

**Fig 4.**
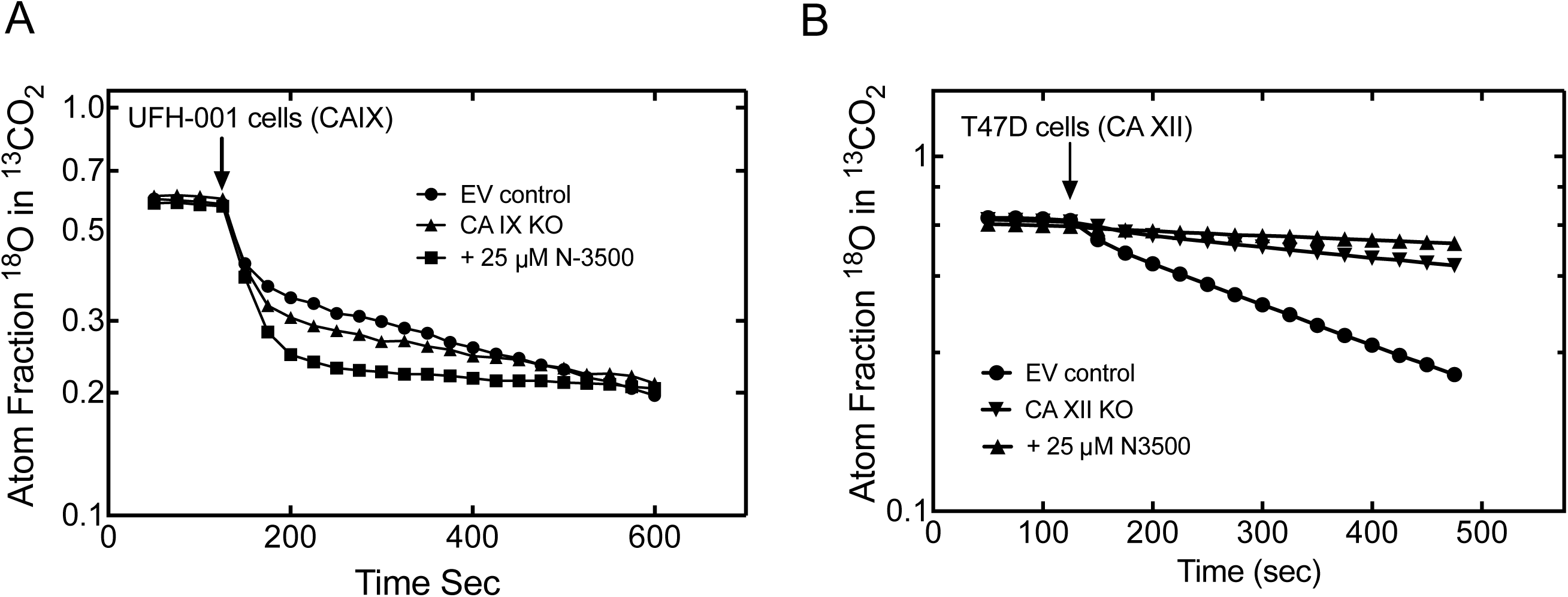
Effect of CA IX and CA XII knockdown on CA activity. CA IX knockout in UFH-001 cells was accomplished using Crispr/Cas9 technology. CA XII knockdown in T47D cells was performed using shRNAi lentiviral strategies. Cells were grown similarly to those described in Fig 3. Panel A. CA activity was measured in 1 x 10^6^ UFH-001 cells (EV controls, CA IX KO, or cells treated with the pegylated sulfonamide, N-3500). Panel B. CA activity was measured in 5 x 10^5^ T47D cells (EV controls, CA XII KO, or cells treated with N-3500. Data represent duplicate experiments.

### Impact of USB-based inhibitors on breast cancer cell growth, cytotoxicity, and apoptosis

MTT, lactate dehydrogenase (LDH), and caspase activity assays were used to test the efficacy of CA IX and CA XII inhibition on breast cancer cell growth, viability and apoptosis, respectively. These results show that all three compounds decreased cell growth (MTT assay) in the different cell lines under normoxic conditions (Fig 5, Table 3). No significant trend in compound effectiveness was established between the cells treated for 48 h compared to 96 h (Fig 5 and Table 3). The most effective compound at inhibiting growth was U-NO_2_ in UFH-001 cells (IC_50_ ∼25 μM). This compound also inhibited the growth of MCF 10A cells in addition to that of the T47D cells, but requiring nearly three times the concentration compared to UFH-001 cells (Table 3). The MCF 10A cells were the most resistant to compounds U-CH_3_ and U-F, with IC_50_ values observed in the range of 500 µM. The concentration required to block growth of T47D cells was also in this range at 48 h but improved somewhat at 96 h of exposure (Table 3). We repeated the experiments under hypoxic conditions (Fig 6) with similar results. In addition, we tested the effects of the USB’s on cells in which CA IX or CA XII was knocked out in UFH-001 (UFH-001 KO) or T47D (T47D KO) cells, respectively, which have been previously characterized [72]. Surprisingly, the result was the same, i.e., the USB’s still blocked cell growth at high concentrations even in the absence of CA IX or CA XII expression.

**Fig 5.**
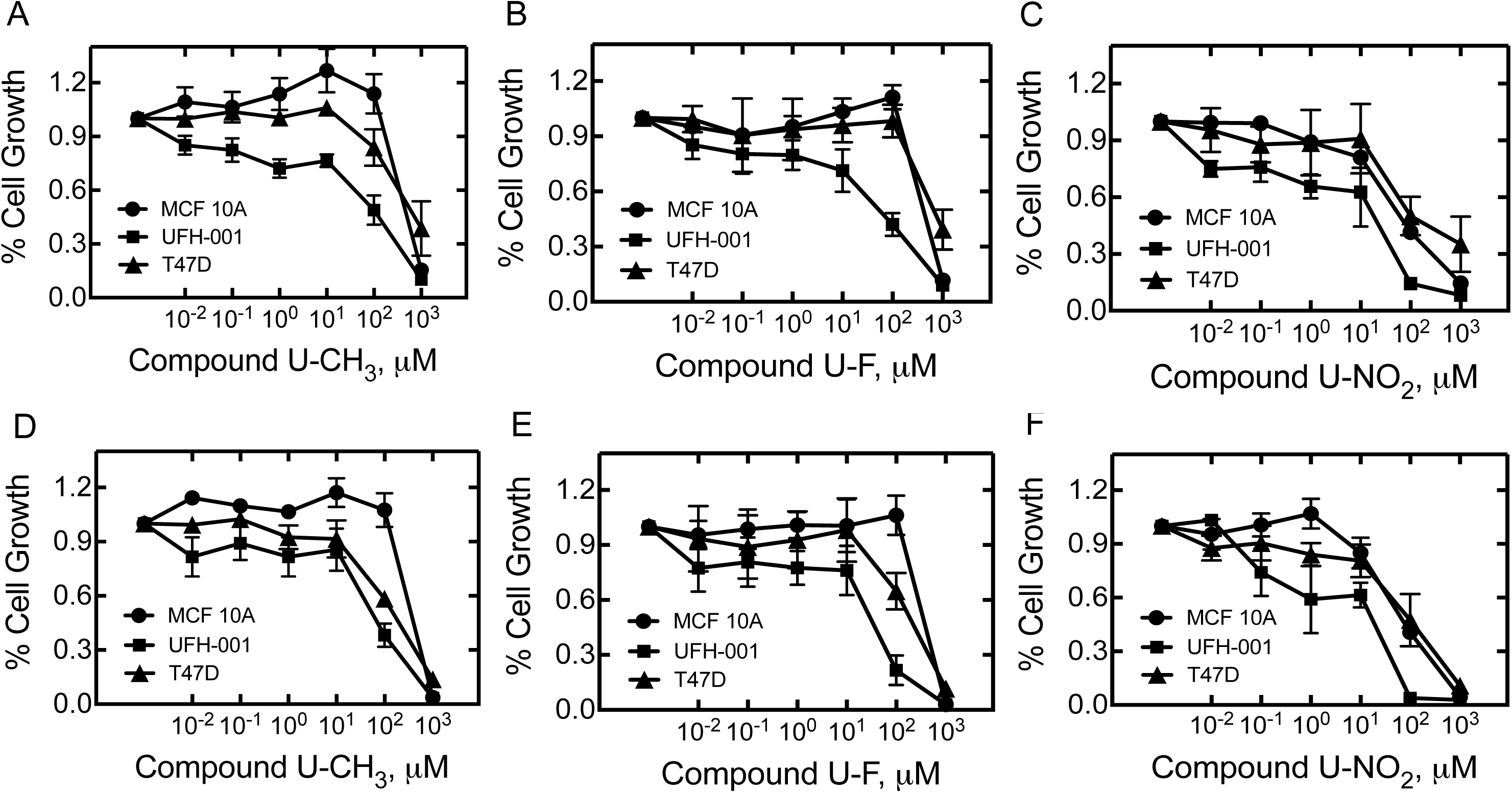
Effects of USBs on breast cancer cell growth. Breast cancer cell lines grown under normal culture conditions for 24 h were exposed to compounds for 48 h [U-CH_3_ (Panel A), U-F (Panel B), U-NO_2_ (Panel D)] or 96 h [U-CH_3_ (Panel D), U-F (Panel E), or U-NO_2_ (Panel F)]. MTT assay was performed at 48 h and 96 h, respectively. Data shown are an average of at least three independent experiments and are represented as the mean ± SEM.

**Fig 6.**
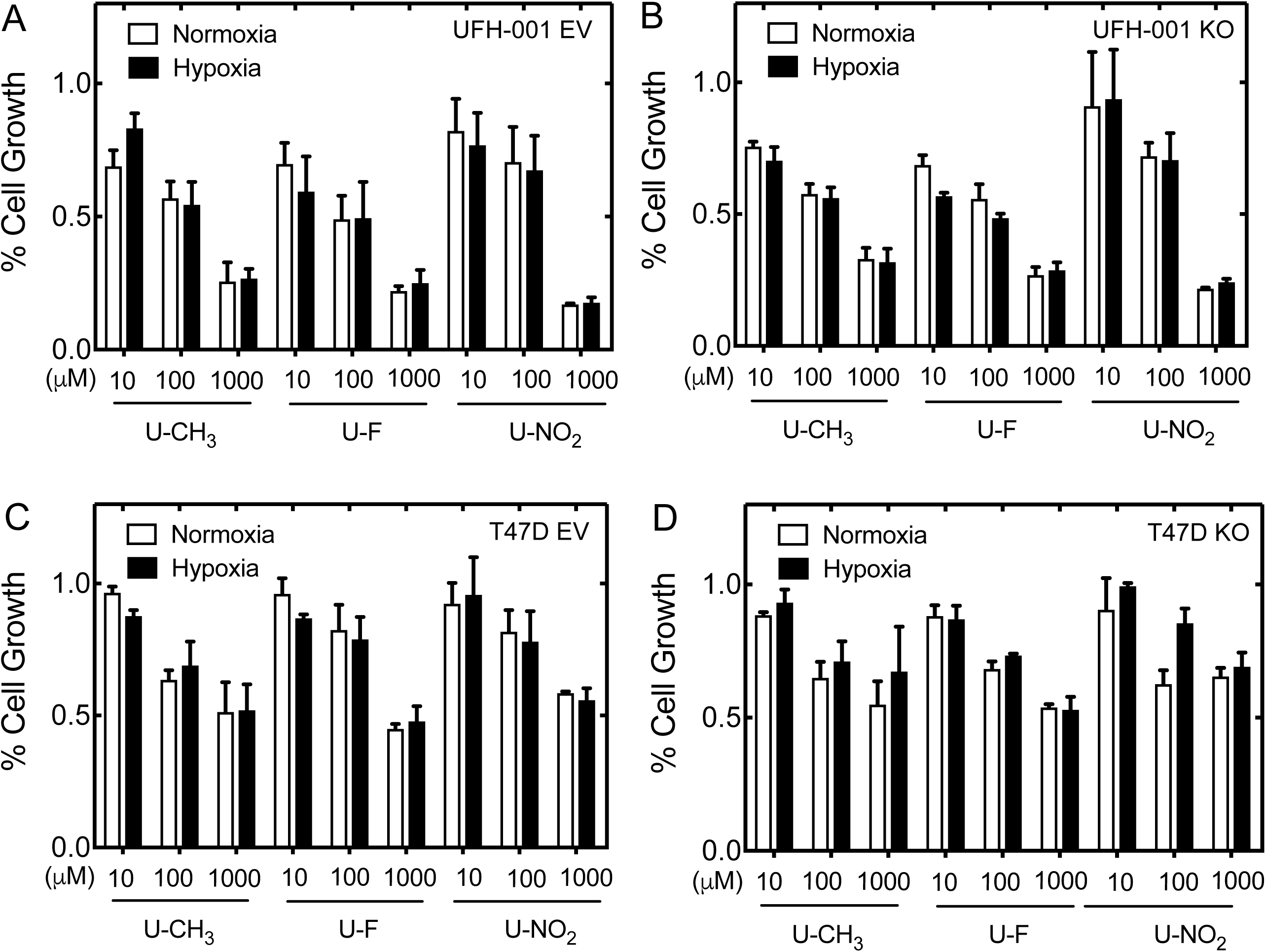
Effects of USBs on cell growth in the presence or absence of hypoxia or in CA knockout cells. One day after plating, breast cancer cell lines were exposed to normoxic or hypoxic conditions for 48 h in the presence of U-CH_3_, U-F, or U-NO_2_ at the given concentrations in UFH-001 empty vector control (EV) cells (Panel A), UFH-001 CA IX KO (UFH-001 KO) cells (Panel B), T47D EV cells (Panel C) and T47D CA XII KO (T47D KO) cells (Panel D). MTT assay was performed and shown as % cell growth. Data shown represent at least three independent experiments and are shown as the mean ± SEM.

**Table 3.** IC_50_ values for compounds **U-CH_3_, U-F,** and **U-NO_2_** in MCF 10A, UFH-001 and T47D cells obtained from MTT assay experiments.

| Cell Lines | MCF 10A Cells |  | UFH-001 Cells |  | T47D Cells |  |
| --- | --- | --- | --- | --- | --- | --- |
| Treatment | 48 h | 96 h | 48 h | 96 h | 48 h | 96 h |
| **U-CH<sub>3</sub> (μM)** | 491 ± 10 | 508 ± 10 | 80 ± 2 | 74 ± 1 | 330 ± 10 | 101 ± 10 |
| **U-F (μM)** | 550 ± 10 | 550 ± 10 | 62 ± 1 | 51 ± 1 | 537 ± 10 | 252 ± 20 |
| **U-NO<sub>2</sub> (μM)** | 64 ± 1 | 69 ± 1 | 26 ± 1 | 20 ± 1 | 61 ± 1 | 78 ± 1 |

Both UFH-001 and T47D cells are able to form spheroids (S4 Fig). UFH-001 cells form spheroids as early as 24 h after plating (even when the plating density is low). The spheroids appear dense and round. The T47D cells also form spheroids, which are somewhat larger than UFH-001 spheroids. Neither the loss of CA IX nor CA XII affects spheroid formation over the 96 h observation period. We were unable to measure activity in these spheroids because of the limited number of cells.

To specifically test for cytotoxicity, LDH release over 48 h in the presence or absence of the USB compounds was measured. The inhibitors caused only limited cytotoxicity when exposed to cells (Fig 7), relative to a positive control (S5 Fig). UFH-001 cells were the most sensitive, but only at high concentrations of compound U-NO_2_ (Fig 7). The MCF 10A cells were also somewhat sensitive to U-NO_2_ but again at high concentrations (Fig 7A). T47D cells showed no loss of membrane permeability in response to inhibitor treatment (Fig 7C).

**Fig 7.**
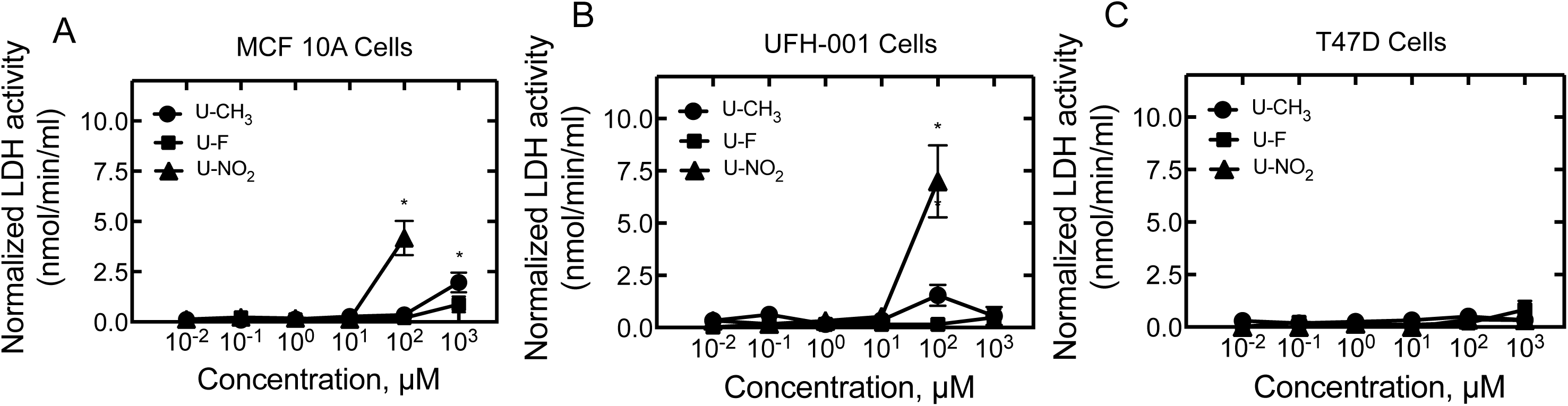
Effects of USBs on breast cancer cell viability. The cytotoxic effects of USBs were evaluated using the LDH release assay. MCF 10A (Panel A), UFH-001 (Panel B) and T47D (Panel C) cells were grown in 96-well plates and exposed to U-CH_3_, U-F or U-NO_2_ for 48 h, under normoxic conditions. LDH release was assayed after treatment, results were evaluated, and data analyzed using Prism. Data shown are representative of three independent experiments and as the mean ± SEM, *p < 0.05.

These observations led us to question whether the growth arrest and (limited) cytotoxicity observed with the inhibitors was due to the activation of apoptosis. To test this, caspase activity assays were employed. These data revealed that apoptosis was not activated, as caspase activity did not change in either UFH-001 or T47D cells in the presence of USB compounds (S6A and S6B Fig, respectively).

Because of the USB-induced growth arrest, albeit at high concentrations, the effects on cell cycle transition were evaluated after exposure to inhibitors for 48 h, under normoxic conditions (Fig 8). Of the diploid population, ∼ 40% of the UFH-001 cells were in the G1 phase of the cell cycle (Dp G1) and a much lower percentage were in G2 phase (Dp G2). The number of T47D cells in G1 (∼ 60%) was significantly higher than the number of cells in the G2 phase of the cell cycle (∼ 20%). A significantly higher percentage of cells were observed in the G1 phase in T47D compared to UFH-001 cells, while cells in G2 phase were much higher in UFH-001 than T47D cells (Fig 8C). In the presence of USB compounds, there was no change in the number of UFH-001 cells in S phase when compared to the control cells (Fig 8D). However, a significant decrease in the number of T47D cells in S phase was noted with exposure to U-CH_3_ and U-F, but not U-NO_2_ (Fig 8E). Higher concentrations of U-CH_3_ and U-F where used for experiments with T47D cells, because both the K_i_ values obtained from the MIMS experiments (Fig 3G) and the IC_50_ values obtained from the cell growth assays (Table 3) were much higher for these compounds relative to compound U-NO_2_, and by comparison to the more sensitive UFH-001 cells.

**Fig 8.**
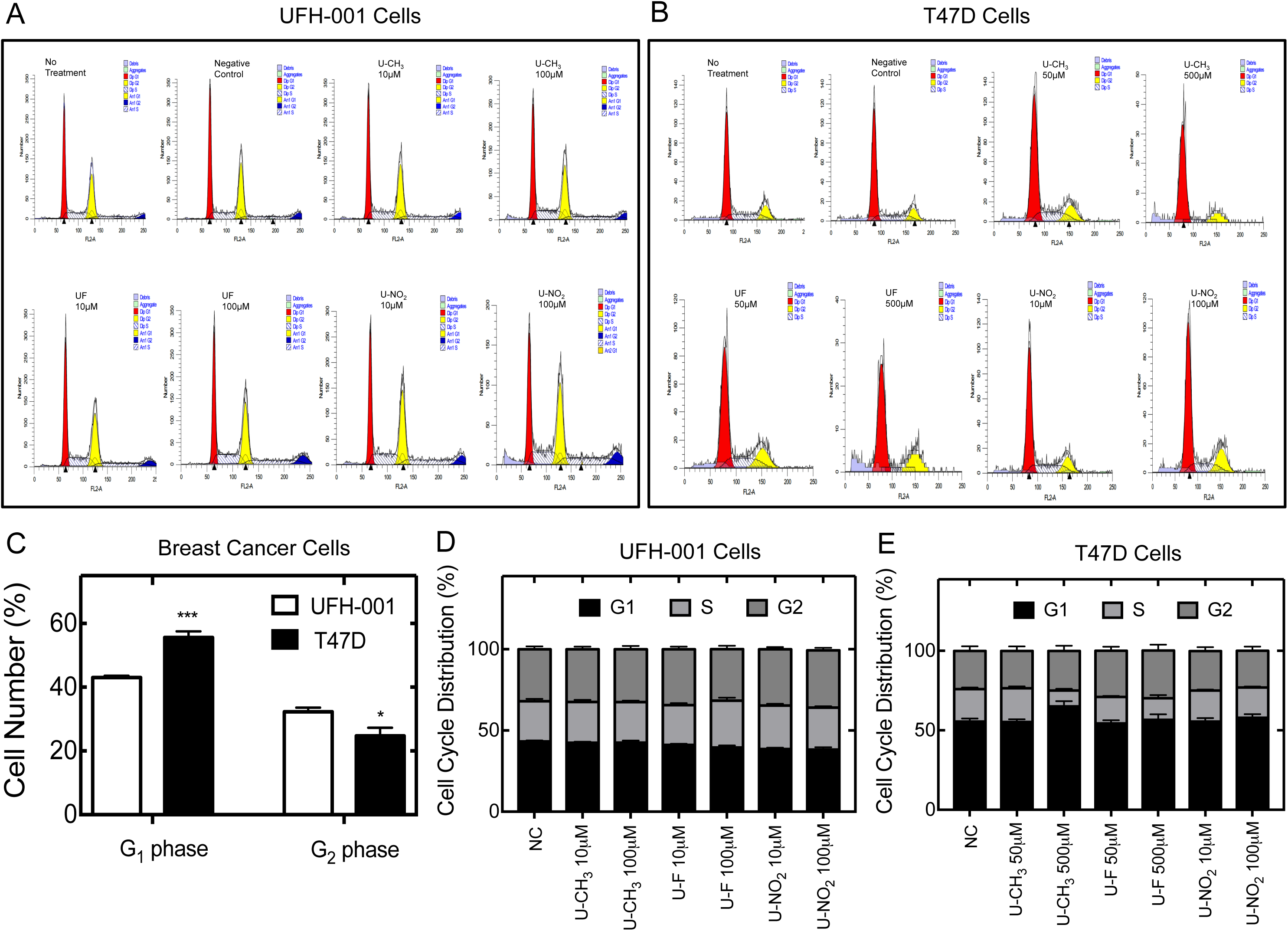
Effects of USBs on cell cycle transition in breast cancer cells. Cell cycle analysis was performed in UFH-001 (Panel A) and T47D (Panel B) cells treated with varying concentrations of USB compounds for 48 h, under normoxic conditions. Post treatment, cells were stained with Propidium Iodine containing RNase A, data was obtained using the FACS caliber instrument and results analyzed with FCS Express and ModFit LT softwares. The percentage of breast cancer cells in G-phase (Panel C), all phases of the cell cycle in UFH-001 (Panel D) and T47D cells (Panel E), were quantified using Prism. Data are represented as mean of at least three independent experiments ± SEM and NC = negative control.

### Effect of USB inhibitors on CA expression in breast cancer cells

We next examined if inhibition of CA IX and CA XII activity induced an upregulation of the targets or expression of other CA isoforms as a compensatory mechanism. To achieve this, UFH-001 and T47D cells were treated with increasing concentrations of USB compounds for 48 h, under normoxic conditions. Immunoblotting with cell lysates was performed using antibodies against CA II, CA IX and CA XII. No upregulation of other CA isoforms was observed in response to USB treatment (S7 Fig). However, there were changes in the endogenously expressed CAs. For instance with U-CH_3_ treatment, both UFH-001 and T47D cells showed increases in the expression of CA IX and CA XII, respectively, compared to controls (S7A and S7B Fig, respectively). Treatment with compound U-F, at lower concentrations (1 μM and 10 μM) increased CA IX and CA XII expression, in UFH-001 and T47D cells, respectively when compared to the control. However, at higher concentrations (100 μM) a decrease in CA IX and CA XII expression was observed (S7A Fig) and in some cases even to a lesser extent than the control cells (S7B Fig). Similar to U-F, compound U-NO_2_ also increased, somewhat, the expression of CA IX and CA XII at lower drug concentrations but ultimately reduced expression, more so with CA IX than CA XII, at concentrations of 100 μM of the compound, when compared to the untreated controls (S7A and S7B Fig).

### Effect of USB inhibitors on cell mobility

While cells grown on polystyrene plates provide useful models, studies of cells grown on more compliant surfaces allow them to recapitulate, in part, their behavior *in vivo*. We have taken advantage of Boyden Chambers to monitor migration and invasion of UFH-001 and T47D cells in the presence or absence of USB compounds. The results show that only the UFH-001 cells have the ability to migrate and invade which was tested at 24 and 48 h, respectively (Fig 9A and 9C). All three USB compounds blocked UFH-001 cell migration at high concentrations: U-CH_3_ and U-NO_2_ were the most effective. Significant inhibition of UFH-001 cell invasion was observed with only U-F treatment and at the highest concentration, well above their K_i_ values for inhibition of activity. Not unexpectedly, we did not observe T47D cell migration and invasion in either the absence or presence of the USBs (Fig 9B and 9D) and thus we are unable to make a definitive statement regarding any potential effect of the sulfonamide inhibitors on cells that do not possess these features.

**Fig 9.**
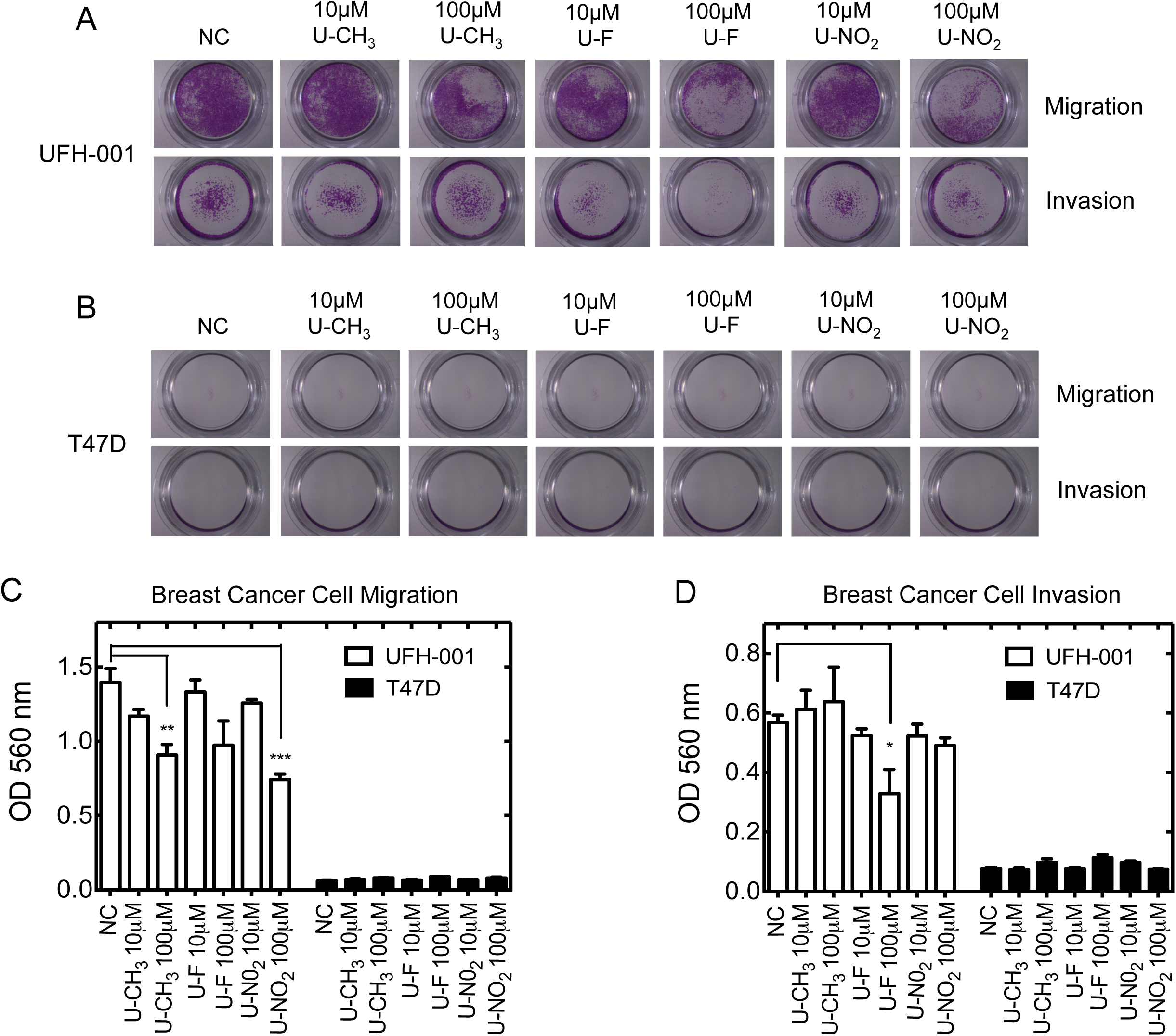
Effects of USBs on breast cancer cell migration and invasion. Serum starved UFH-001 and T47D cells were allowed to either migrate for 24 h or invade for 48 h towards a chemoattractant in the presence or absence of USB compounds. Bright-field images of migrating and invading cells are shown in Panel A (UFH-001 cells) and Panel B (T47D cells). Data are quantified for UFH-001 and T47D cells Panel C (migration) and Panel D (invasion). NC= negative control. Data shown are an average of duplicate experiments ± SEM. *p < 0.05, and ***p < 0.001.

## Discussion

In the current study, we investigated the binding of a class of USB-based compounds that were specifically designed to inhibit the cancer-associated isoforms, CA IX and CA XII, over the off-target CA II. We have shown, as others have previously, that residue 131 is important for isoform selectivity of these compounds towards CA IX and CA XII [66-68, 77, 83]. In addition, our model shows that serine residues 132 and 135 located within the active site of CA XII are required for USB selectivity (compared to the other CAs), towards this isoform. This then raises the question: Can we relate these structural data to the K_i_ values published for recombinant CAs [68]? Data in Table 1 show that compounds U-CH_3_ and U-F exhibit K_i_ values for CA II that are one to two orders of magnitude greater than for CA IX or CA XII. Our structural data illustrate that these differences are caused by residue 131, which is a phenylalanine in CA II, forcing a lower energy conformation of U-CH_3_ and U-F in the active site of CA II [66]. This differs from the more favorable conformation adopted by inhibitors in CA IX [66] and predicted in CA XII (Fig 1). U-NO_2_ conformation is similar among the three CA isoforms, hence the similar K_i_ values. Therefore, we must conclude that residue 131 does not cause steric hindrance in the active site of CA II for U-NO_2_ but clearly does create steric hindrance for the other two inhibitors. This may be related to physiochemical properties or flexibility of the tail moiety.

In the physiological setting of intact cells, inhibition of CA IX activity in UFH-001 cells by USBs was significantly more efficient than inhibition of CA XII in T47D cells (Fig 3). This was unexpected based on the relatively similar K_i_ values obtained using recombinant proteins (Table 1) [68]. This difference in efficacy was about an order of magnitude between the cell line that expresses CA IX versus the cell line that expresses CA XII. Because of the difference in CA IX and CA XII as prognosticators for patient outcome, this result suggests the potential for selective therapeutic targeting for CA IX, in the clinical setting, despite the similar efficacy of the drugs with the purified proteins. However, there was also a difference in K_i_ values between purified protein and those in intact cells. For CA IX, this difference varied across inhibitors ranging from 3-fold (U-F) to 500-fold higher (U-NO_2_) (i.e., inhibition of CA IX in cells versus recombinant protein). For CA XII, this difference was nearly three orders of magnitude for all USBs. It is possible that part of this difference lies in the use of different techniques for analyzing activity. Recombinant protein activity was measured using stop flow kinetics [55] while enzymatic activity in cells was measured using MIMS, a technique originally developed to define the catalytic mechanisms of purified CAs [75, 82], including that for CA IX [84]. That differences in the K_i_ values are based on technique seems unlikely because the rate constants calculated for recombinant CA IX are similar across these techniques [84-86]. Further, rate constants determined in isolated membranes containing either CA IX or CA XII [27, 72] show strikingly similar values, suggesting that the membrane environment does not influence catalytic activity. Yet, we must also consider that the cellular environment (intact cells) adds layers of complexity relative to even isolated membranes. The cells are in suspension during the MIMS assay, having been released from plates using a non-enzymatic procedure. Thus, many of the cells are still in “colonies” which means that the extracellular matrix is still intact. This layer, comprised of glycoproteins, proteoglycans, and fibrous proteins like collagen, may provide a barrier for incoming molecules (like USBs). Yet, the permeant sulfonamide, ethoxzolamide, inhibits activity even inside the cell and with no time delay. In addition, the USBs are impermeant over the course of the assay, and thus inhibitor concentration is preserved outside of the cells.

The effect of the USB inhibitors on cell growth followed a trend similar to that observed for activity, but is more problematic. For both the UFH-001 and T47D cells, the inhibitors were less effective by about two orders of magnitude, based on IC_50_ values. We initially conducted these experiments under normoxic conditions, even though aggressive forms of cancer exist in hypoxic environments. Our rationale for using normoxia was based on the lack of difference in the K_i_ values of the USB compounds for CA IX and CA XII between normoxic and hypoxic cell culture conditions. However, Elena et al. showed that U-F (SLC-0111) activated apoptotic and necrotic programs only in acidified medium [87], a condition associated with hypoxia. But again, these effects were seen at only high concentrations of inhibitor, orders of magnitude above the K_i_ values that we have measured for inhibition of CA activity. Regardless, we repeated the growth studies with cells exposed to hypoxia, but the data were identical. Several studies have shown that these USB compounds are effective at blocking tumor growth [68, 88, 89], so this is clearly a useful strategy for treating breast cancer patients. So, we must ask why there is a disconnect between the loss of CA activity, particularly that of CA IX, and the inhibition of cell growth/cytotoxicity. The activity assay is conducted for about 6 min, while the growth, migration/invasion, and cytotoxicity experiments are conducted for up to 96 h. Under these later conditions it is possible that the inhibitors undergo chemical inactivation by oxidation, reduction, or cleavage reducing their ability to block the CAs. It is difficult to measure these parameters in the context of cell culture. It is also possible that the drugs are transported into cells over the long term, reducing the local concentration around exofacial catalytic sites of CA IX or CA XII. However, the studies in which we tested USB compounds, in cells where CA IX and CA XII were ablated, showed the same inhibitory effect of sulfonamides on cell growth. This suggests that, at very high concentrations of the USB’s, they are acting independently of their inhibition of exofacial CA activity. Indeed, others have recently observed this as well [90, 91]. In these latter studies, investigators identified a potential new sulfonamide target, RMB39 (RNA binding motif protein 39) that is also called CAPERα. They demonstrated that indisulam (an aryl sulfonamide drug in phase II clinical trials for the treatment of advanced stage solid tumors) promotes the recruitment of RBM39 to the CUL4-DCAF15 E3 ubiquitin ligase leading to RBM39 ubiquitination and proteasomal degradation [90, 91]. Mutations in RBM39 that prevent its recruitment increase its stability, conferring resistance to indisulam’s inhibition of cell growth and cytotoxicity [90]. Another series of studies used a naphthalene sulfonamide to bind to the pY705 in STAT3 (signal transducer and activator of transcription 3). This binding inhibits STAT3 phosphorylation and dimerization, which is required for its interaction with DNA [92, 93]. *In vivo*, this sulfonamide blocks tumor growth in a breast cancer xenograft. With these “off target” effects of sulfonamides, we must consider that exofacial CA activity, alone, does not regulate cell growth.

In conclusion, we have shown that specific residues within the catalytic sites of CA IX and CA XII determine the binding affinities of the USB inhibitors. These interactions, particularly at residue 131, provide the rationale for selective inhibition of recombinant CA IX and CA XII over the off-target CA II. Based on K_i_ values, determined in the context of cells, we conclude that USB inhibition of activity shows strong specificity for CA IX relative to CA XII. This was surprising based on the similar K_i_ values between recombinant CA IX and CA XII. Because there are now data to further prove that CA IX acts to stabilize pH in the tumor setting, blocking its activity may alter the microenvironment, specifically in those tumors that express CA IX over CA XII. While we observe the same trend in cell growth inhibition, (i.e., UFH-001 cells are more sensitive to USB inhibition than T47D cells), the inhibitor IC_50_ values increase by another two orders of magnitude. This leads us to question the role of CA activity in cell growth. Indeed, the presence of CA IX or CA XII is not required for USB-mediated cell growth inhibition. Obviously, the inhibitors do block cell growth in culture and tumor growth in xenograft and metastatic models, so their value as chemotherapeutic agents may still be considerable. That said, the mechanism of action of sulfonamides need further investigation.

## Acknowledgements

The authors would like to recognize the exceptional cell culture skills of Xiao Wei Gu.

## Supporting information

**S1 Fig. Effect of classical sulfonamide inhibitors on CA activity in UFH-001 and T47D cells.** CA activity was measured in UFH-001 cells (Panel A) and T46D cells (Panel B) using the MIMS assay in the absence or presence of acetazolamide (ACZ) or ethoxzolamide (EZA). Data are representative of two independent experiments. First order rate constants were calculated according to the formula described in the Methods. Is noted that the scale on the y-axis is different between these two representative plots. This difference represents the different isotopic enrichments of CO_2_, but the concentration of CO_2_ is identical between the two experiments.

**S2 Fig. CA mRNA expression in breast cells lines.** Panel A: mRNA expression (from microarray data) in a normal immortalized basal type breast cell line (MCF10A) compared to a triple negative breast cancer cell lline (UFH-001) and Panel B: MCF10A versus T47D cells were analyzed using data mining techniques. Accession numbers, GSE107209 (for comparison between MCF10A and UFH-001 cell lines) and NCI-60 data sets for T47D cells, were used for this comparison.

**S3 Fig. Effect of U-NO_2_ on CA activity.** CA activity was measured in normoxic and hypoxic cells UFH-001 cells (Panel A) or normoxic and hypoxic T47D cells (Panel B) in the presence of U-NO_2_ to determine K_i_ values across an extensive range of inhibitor concentrations.

**S4 Fig. Effect of CA knockdown on spheroid growth.** Western blots of lysates from UFH-001 cells (EV controls and KO cells) exposed to normoxic or hypoxic conditions (Panel A) were compared to lysates from T47D cells (EV controls and KO cells) exposed to normoxic and hypoxic conditions (Panel B). Panel C shows spheroid development of UFH-OO1 cells (EV controls and KO cells) while Panel D shows spheroid development of T47D cells (EV controls and KO cells) over 96 h in culture. GAPDH and actin were used as loading controls.

**S5 Fig. Total LDH activity released by breast cell lines.** Cells were grown in 96 well plates for 24 h at which point they were treated with a drug which is cytotoxic (β-caryophyllene) as a positive control or left untreated (NC) under normoxic conditions. LDH assays were performed after 48 h of treatment, results were evaluated at 450 nm (absorbance), and data was analyzed using Prism. Total LDH activity (nmol/min) ws assessed in Panel A) MCF10A cells; Panel B UFH-001 cells; and Panel C t47D cells. Data represent the mean ± SEM of 3 independent experiments.

**S6 Fig. Effect of USBs on activation of apoptosis.** Activation of apoptotic pathways was evaluated using the caspase activity assay in Panel A) UFH-001 and Panel B) T47D cells after 48 h of treatment with either absence (negative control, NC) or presence of USB-based compounds, under normoxic conditions. These data were compared to the presence of staurosporine (positive control, PC). Data shown for the USB-treated cells are the averages of at least three independent experiments. For the PC-treated cells, these data represent the average of two independent experiments.

**S7 Fig. Effects of USB compounds on CA expression in breast cancer cells.** Immunoblotting is shown for CA IX, CA XII, and CA II from cells grown for 2-3 days and then treated with compounds U-CH3, U-F, or U-NO2 for 48 h, under normoxic conditions. GAPDH was used as a loading control. Data are representative of 3 independent experiments. Panel A, UFH-001 cells. Panel B, T47D cells.

